# GO: A functional reporter system to identify and enrich base editing activity

**DOI:** 10.1101/862458

**Authors:** Alyna Katti, Miguel Foronda, Jill Zimmerman, Bianca Diaz, Maria Paz Zafra, Sukanya Goswami, Lukas E Dow

## Abstract

Base editing (BE) is a powerful tool for engineering single nucleotide variants (SNVs) and has been used to create targeted mutations in cell lines, organoids, and animal models. Recent development of new BE enzymes has provided an extensive toolkit for genome modification; however, identifying and isolating edited cells for analysis has proven challenging. Here we report a “Gene On” (GO) reporter system that indicates precise cytosine or adenine base editing *in situ* with high sensitivity and specificity. We test GO using an activatable GFP and use it to measure the kinetics, efficiency, and PAM specificity of a range of new BE variants. Further, GO is flexible and can be easily adapted to induce expression of numerous genetically encoded markers, antibiotic resistance genes, or enzymes such as Cre recombinase. With these tools, GO can be exploited to functionally link BE events at endogenous genomic loci to cellular enzymatic activities in human and mouse cell lines and organoids. Thus, GO provides a powerful approach to increase the practicality and feasibility of implementing CRISPR BE in biomedical research.

---

Base editing (BE) is a powerful genome engineering tool that harnesses Cas9-mediated gene targeting to induce specific point mutations in DNA or RNA (1). Base editors consist of: 1) a partially enzymatically disabled Cas9 protein (Cas9n, or ‘nickase’) to enable genomic targeting, 2) a fused nucleobase deaminase to catalyze transition mutations, and in some cases, 3) one or more uracil glycosylase inhibitor (UGI) domains, which enhance base conversion by mitigating endogenous DNA repair activity (2). Cytosine base editors (CBEs) use APOBEC or AID deaminase domains to induce C>T (or G>A) mutations (3, 4) while adenine base editors (ABEs), use an engineered bacterial protein, TadA, to introduce A>G (or T>C) changes (5). In theory, more than half of all pathogenic point mutations can be introduced or reversed by BE (2, 5–7), making it a powerful approach to model disease-associated single nucleotide variants (SNVs). Indeed, numerous studies have highlighted the power of BE to engineer defined alterations in cell lines, organoids, and *in vivo* in a diverse array of model systems (2, 3, 6, 8–11).

Unlike Cas9, which shows remarkable efficacy in creating homozygous disruptive mutations following DNA doublestrand breaks (DSBs), BE is relatively inefficient. BE activity depends on the level of enzyme expression, sequence context of the target site, cellular DNA repair, and likely other unexplained dependencies. Identifying and enriching BE events in cells is critical to streamline the use of these tools for biomedical research. To date, a number of BE reporters have been described, each using distinct mechanisms, but all ultimately based on the induction or suppression of GFP fluorescence. Hence, identification of base edited, live cells by BE reporters has thus far relied on fluorescence-based imaging or cell sorting (FACS) of a single fluorophore, somewhat limiting their broad application across different cell systems.

Here, we describe a flexible “Gene On” (GO) functional reporter system that enables detection and enrichment of BE activity in living cells. We show that GO can be used to directly and quantitatively compare the efficiency, off-target activity, and PAM selectivity of existing and novel BE enzymes. Most importantly, because GO is based on translation initiation, it is not limited to the regulation of GFP, but enables the induction of different fluorescent and bioluminescent markers, antibiotic selection, or functional enzymes such as Cre-recombinase. Thus, GO is a specific, flexible and functional BE reporter that can identify and enhance the application of base editing activity in primary and immortalized cell lines and organoids.

## RESULTS

### Development of a specific cytosine base editing reporter: Cy^GO^

To detect DNA cytosine BE events in real time in living cells, we sought to create a reporter that would become activated upon APOBEC-mediated C>T conversion, but not following Cas9-mediated indels. Ribosomal protein translation requires an initiating AUG codon immediately downstream of a Kozak sequence (e.g. gccRccAUGG) in messenger RNA (mRNA) transcripts. Therefore, we hypothesized that the de novo creation of an ATG codon (from ACG) at the start of a cDNA would enable translational initiation and production of a specific protein detectable in individual cells with active cytosine BE (Cytosine Gene On, or “Cy^GO^”; Figure 1a).

**Figure 1.**
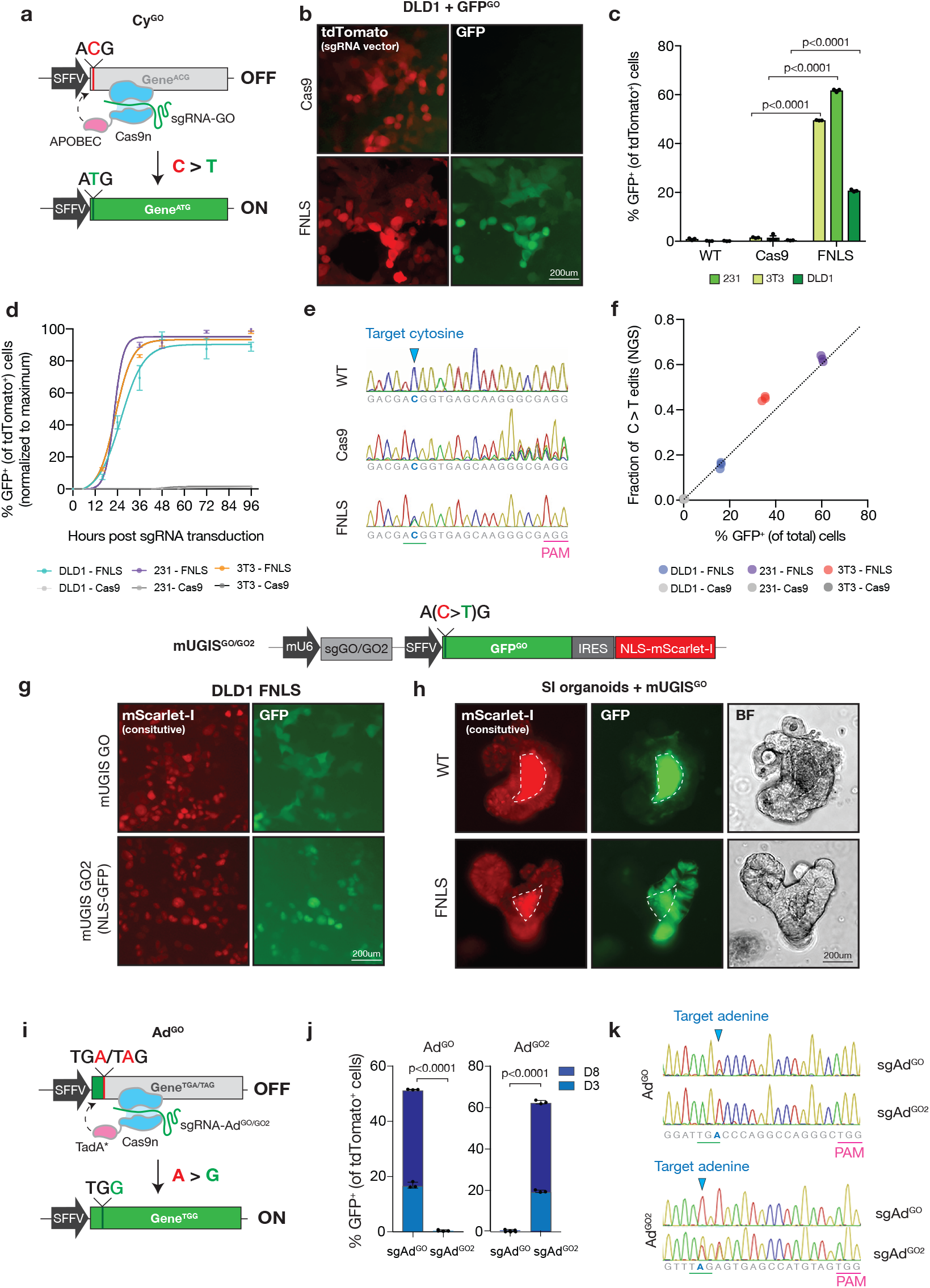
GO detects and reports cytosine and adenine base editing with high sensitivity and specificity. **a.** Schematic showing cytosine BE (Cy^GO^) system containing 1) a cDNA with a silent ‘ACG’ start site 2) FNLS base editor, and 3) sgRNA targeting the 5’‘ACG’ for cytosine deamination. **b.** Fluorescence imaging of DLD1s transduced with GFP^GO^, sgGO-tdTomato, and Cas9 or FNLS. **c.** Flow cytometry analysis of DLD1, 231, and 3T3 cells transduced with GFP^GO^ and Cas9, FNLS, or no editor (WT) 3 days after sgGO-tdTomato infection. GFP^+^ cells were gated on tdTomato^+^ cells. **d**. Time course flow cytometry of GFP induction after sgGO-tdTomato transduction. **e.** Sanger sequencing chromatograms of GFP^GO^ locus in 3T3s containing GFP^GO^ and Cas9, FNLS, or no editor (WT) 3 days post sgGO-tdTomato infection. **f.** Correlation of C>T editing with percentage of GFP^+^ cells (gated on total cells). **g.** Schematic displaying an all-in-one Cy^GO^ containing 1) sgGO, 2) GFP ‘ACG’ gene, and 3) constitutive marker mScarlet-I (mUGIS^GO^). Including a GFP tagging 5’ nuclear localization signal and modified 5’ targeting sgRNA (sgGO2) generated mUGIS^GO2^. Fluorescence microscopy of DLD1s expressing FNLS 3 days post infection with mUGIS^GO^ or mUGIS^GO2^. **h.** Fluorescence and brightfield imaging of small intestinal murine organoids infected with mUGIS^GO^ 3 days after FNLS or no editor (WT) transfection. Dashed lines indicate auto-fluorescent lumen. **i.** Schematic of adenine BE (Ad^GO^) system containing 1) a cDNA with a premature ‘TAG’ or ‘TGA’ stop codon, 2) ABE base editor, and 3) sgRNA targeting the 5’ cDNA at the early stop codon (sgAd^GO^). Ad^GO2^ and sgAd^GO2^ were designed with a modified 5’ region. **j.** Flow cytometry analysis of 231s expressing ABE and Ad^GO^ or Ad^GO2^ quantifying GFP activation 3- and 8-days post sgAd^GO^-tdTomato or sgAd^GO2^-tdTomato infection. GFP^+^ cells were gated on tdTomato^+^ cells. **k.** Sanger sequencing traces of Ad^GO^ and Ad^GO2^ loci from 231s expressing ABE and Ad^GO^ or Ad^GO2^ 6 days after sgAd^GO^-tdTomato or sgAd^GO2^-tdTomato infection.

To test this idea, we generated a ‘silent’ GFP^ACG^ construct and expressed it in human and mouse cells via lentiviral delivery; as expected, cells transduced with a SFFV-GFP^ACG^-PGK-NeoR (GFP^ACG^) lentivirus showed no detectable GFP fluorescence (Supplementary Figure 1a). We then introduced an sgRNA targeting the 5’ region of GFP^ACG^ (sgGO) in a tdTomato-P2A-BlasR lentivirus (6) into GFP^ACG^ cells expressing either Cas9 or an optimized cytidine base editor (FNLS) (together, referred to as (GFP^GO^)(6). While Cas9/sgGO efficiently targeted the GFP^ACG^cDNA, inducing indels in more than 60% of cases, we did not detect GFP fluorescence above background levels (Figure 1b-c, Supplementary Figure 1b). In contrast, FNLS-expressing cells showed a robust, cell line dependent, induction of GFP fluorescence in immortalized mouse fibroblasts, NIH3T3s (3T3s), and human cancer cell lines, DLD1 and MDA-MB-231 (231s) (Figure 1b-c, Supplementary Figure 1c). In each cell line, editing occurred rapidly, approaching saturation within the first 48 hours following sgGO transduction (Figure 1d). Targeted sequencing revealed a strong linear correlation of C>T (ACG>ATG) base substitutions with the percentage of GFP positive cells (Figure 1e, f), confirming the direct relationship between target BE and GFP signal. As expected, there was no positive association of Cas9 or CBE-induced indels with GFP fluorescence (Figure 1f, Supplementary Figure 1b).

Editing detection was not limited to a single GFP^GO^ reporter or guide design. Changing the GO targeting sequence (sgGO2) enabled induction of an N-terminal nuclear localization sequence (NLS)-tagged GFP^ACG^ cDNA (together, GFP^GO2^) (Supplementary Figure 1d). NLS-GFP expression allows for simple quantification of edited cells by image-based detection (Supplementary Figure 1e). Finally, to support streamlined delivery of the reporter components, we generated an all-in-one vector containing sgGO or sgGO2 under a mouse-human chimeric U6 promoter, latent GFP^GO^ or GFP^GO2^ cDNA, and constitutively expressed marker (i.e. mScarlet-I) to identify transduced cells (mU6-GFP^ACG^-IRES-mScarlet-I with GO or GO2, “mUGIS^GO/GO2^”; Figure 1g). As described in the GFP^GO/GO2^ ‘two-vector’ systems, introduction of the all-in-one reporter mUGIS^GO^ or mUGIS^GO2^ in FNLS-expressing cells showed rapid induction of GFP fluorescence (40-60%) not seen in cells with Cas9 or no editor (Figure 1g, Supplementary Figure 2a-b). As a final validation of the reporter system, we introduced the mUGIS^GO^ reporter into mouse intestinal organoids and observed GFP activation in mScarlet-I-positive cells only after transient transfection with FNLS (Figure 1h). Together, these data show that Cy^GO^ reliably and quantitively detects CBE-mediated C>T transitions but not Cas9-based indels in multiple living cell systems.

### Adenine base editing activates modified GFP^GO^: Ad^GO^

Adenine base editors catalyze the deamination of adenine to inosine, which is recognized as guanine during DNA replication, thus generating A>G transitions. We tested the flexibility of the GO reporter in detecting adenine base editing. We engineered a stop codon, TGA, at the 5’ end of a GFP cDNA (Ad^GO^) to prevent translation extension, and a corresponding sgRNA targeting the 5’ region of GFP^TGA^ (sgAd^GO^) (Figure 1i). We hypothesized that a A>G base substitution by an adenosine base editor (ABE) at this site, making a TGG (Trp) codon, would prevent early protein truncation and promote GFP expression. In 231 cells stably expressing ABE, sgAd^GO^, and Ad^GO^, we detected robust GFP activation (Figure 1j, Supplementary Figure 2c-d). Without ABE, GFP was undetected (Supplementary Figure 2c-d). Again, GFP activation by adenine base editing was not limited to a single reporter/sgRNA combination as GFP activation was detected in a parallel reporter with a modified target stop codon (Ad^GO2^) and sgRNA (sgAd^GO2^) (Figure 1i). Ad^GO^ and Ad^GO2^ reporters were guide specific in that only their respective sgRNA resulted in GFP expression (Figure 1j) which corresponded to A>G base substitutions at the respective Ad^GO/GO2^ locus (Figure 1k). Together, these data show that the GO reporter system can precisely detect both cytosine and adenine base editing in living cells.

### GO enables a quantitative comparison of PAM flexibility and fidelity by BE enzymes

To date, efforts to measure PAM preferences of variant BE enzymes have relied on changing guide target sequences to accommodate the location of a given endogenous PAM site. This may introduce guide dependent effects on editing efficiencies when comparing PAM-flexible editors. To provide a system to directly compare editing efficiency with different PAM sites using the same sgRNA, we engineered four versions of the GFP^GO2^ reporter containing distinct PAM sites (NGG, NGA, NGT, NGC) (Figure 2a). We introduced the four GFP^GO2^ PAM reporters into 231, DLD1, and 3T3 cell lines expressing a series of codon-optimized BE enzymes containing Cas9 variants that have been reported to recognize NGN PAMs and targeted GFP activation by infection with sgGO2 (Figure 2b) (12, 13). As expected, FNLS (containing Cas9n) showed efficient editing of targets with NGG PAMs in all cell lines, but minimal editing of other targets, including NGA (Figure 2c). Similarly, xFNLS (containing xCas9 (12)) showed equivalent editing at NGG targets in 2/3 cell lines, and increased modification of NGT targets in the same cells, though not in 3T3s. FNLS-NG (containing Cas9-NG (13)) had the broadest editing scope, showing equivalent editing at NGG, NGT and NGC sites (Figure 2c). Surprisingly, none of the enzymes displayed substantial editing of targets with NGA PAM sites (Figure 2c). The relative efficiency of FNLS-NG was confirmed at an endogenous genomic locus for two overlapping sgRNAs that share same target cytosine with NGG and NGT PAMs (Figure 2d).

**Figure 2.**
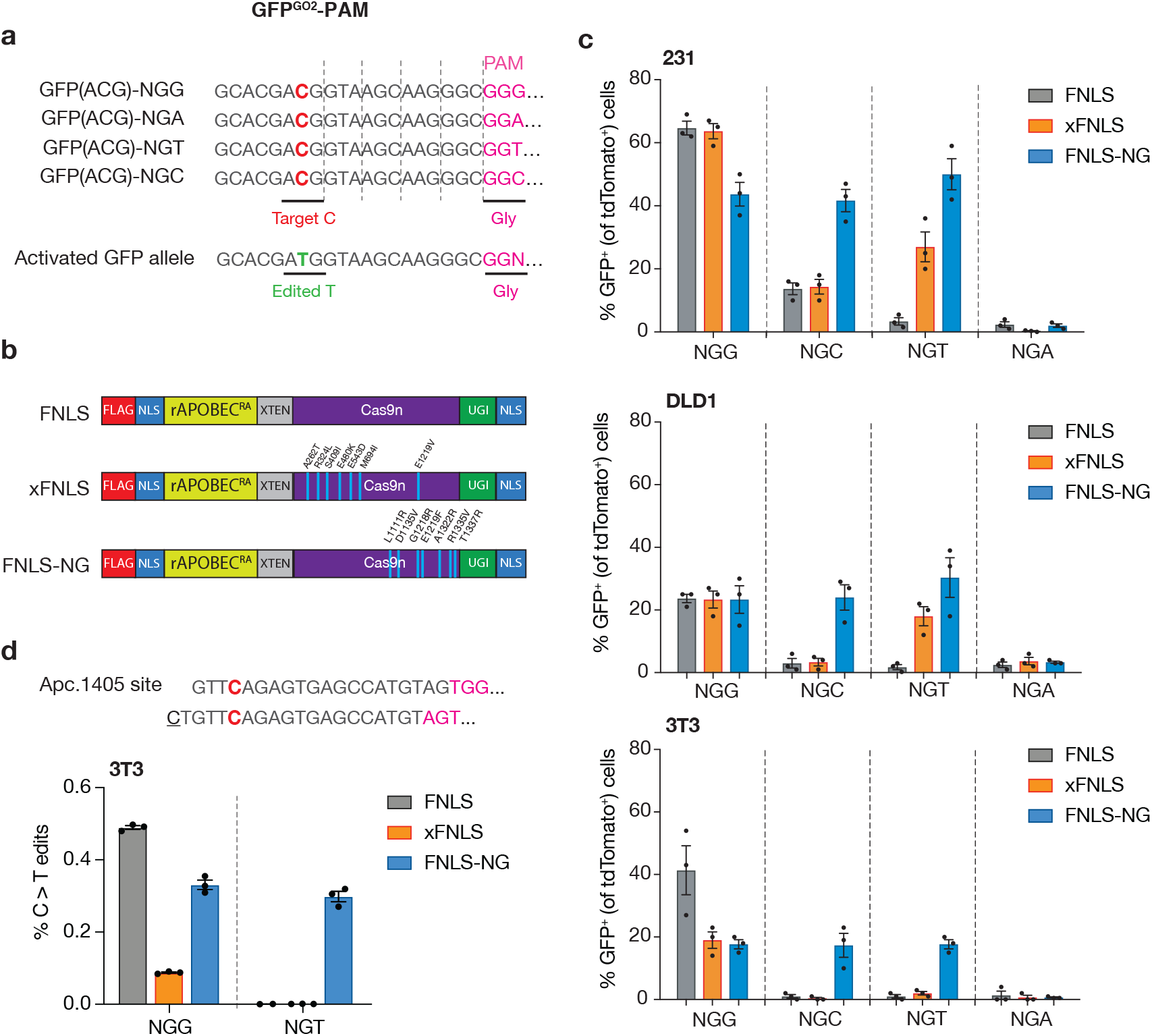
GO quantifies base editing activities of BE enzymes with various PAM specificities. **a.** Panel of GFP^GO^ reporters with four different PAM recognition sequences: NGG, NGA, NGT, NGC designed with constant 20bp complementary target sequence at 5’ end of GFP cDNA for targeting by sgGO-tdTomato. **b.** Schematic of BE PAM variant enzymes included in panel. **c.** Flow cytometry analysis of GFP activation in DLD1, 231, and 3T3 cell lines expressing the panel of editors (b) and infected with each of the four PAM GFP^GO^ reporters 6 days after transduction with sgGO (notated by GFP^GO^ PAM site). GFP^+^ cells were gated on tdTomato^+^ population. **d.** (top) Schematic showing the endogenous Apc locus (codon 1405) in mouse 3T3 cells that contains a cytosine targetable by two nearly identical sgRNA with adjacent and distinct PAM recognition sequences, NGG and NGT. 3T3 cell lines expressing the panel of editors (b) were transduced with either of two nearly identical sgRNAs (NGG or NGT). Deep sequencing analysis of C>T editing events as a fraction of total reads at the Apc.1405 locus in the targeted panel of cells (bottom).

We also demonstrated the utility of GO in quantitatively comparing the fidelity of BE enzyme variants, showing improved specificity of HiFi-FNLS and HF1-FNLS enzymes relative to FNLS (Supplementary Figure 3a-d)(14–16). In addition, we generated a novel combination variant, HiFi-FNLS-NG, that combines high specificity with PAM flexibility at both reporter and endogenous targets (Supplementary Figure 3b-d). We note that individual BE enzymes do not always behave the same in different cell lines, with regards PAM specificity and off-target activity (Supplementary Figure 3c-d). The reason/s for this are unknown, but we expect that the ease of direct BE measurement with flexible reporters such as this, without the requirement for targeted deep sequencing, will provide a feasible setting in which to explore the factors that govern BE activity and fidelity across cell systems.

### GO activates a variety of genetically encoded reporters

As translation initiation at ATG (or less commonly, CTG) is a universal feature of protein coding genes, induction by GO should be generally applicable to almost all genetically encoded reporters. To test the flexibility of GO in reporting BE by a fluorescent protein other than GFP, we generated a ‘silent’ mScarlet-I^ACG^ reporter (Figure 3a) using the same sgRNA sequence for targeting GFP^GO^. We generated an allin-one vector containing sgGO, the mScarlet-I cDNA with 5’ 20bp complementary sequence to sgGO containing the silent ‘ACG’ start site, and a constitutively expressed neomycin resistance gene (Scar^GO^). As described for GFP, mScarlet-I expression was not detectable in WT 231 cells, and was efficiently induced in G418-resistant cells expressing FNLS (Figure 3a). FNLS expressing-3T3s, DLD1s, and 231s cells showed similar base editing efficiencies with Scar^GO^ to the GFP^GO^ reporters (Figure 3c, Supplementary Figure 4a-b). We next tested GO induction of a non-fluorescent marker by replacing the 5’ coding sequence of Luciferase 2 (Luc2) with the sgGO target sequence and silent ‘ACG’ start site (Figure 3b). Transduction of both human 231 cells and immortalized murine embryonic fibroblasts (MEFs) revealed a dramatic induction of Luciferase activity only in cells expressing FNLS (Figure 3b). Thus, GO can be adapted to a variety of genetically encoded reporter constructs, providing a flexible option for identification of activity BE cells in different cell systems and induction of a bioluminescent reporter that will enable imaging of in vivo BE activity.

**Figure 3.**
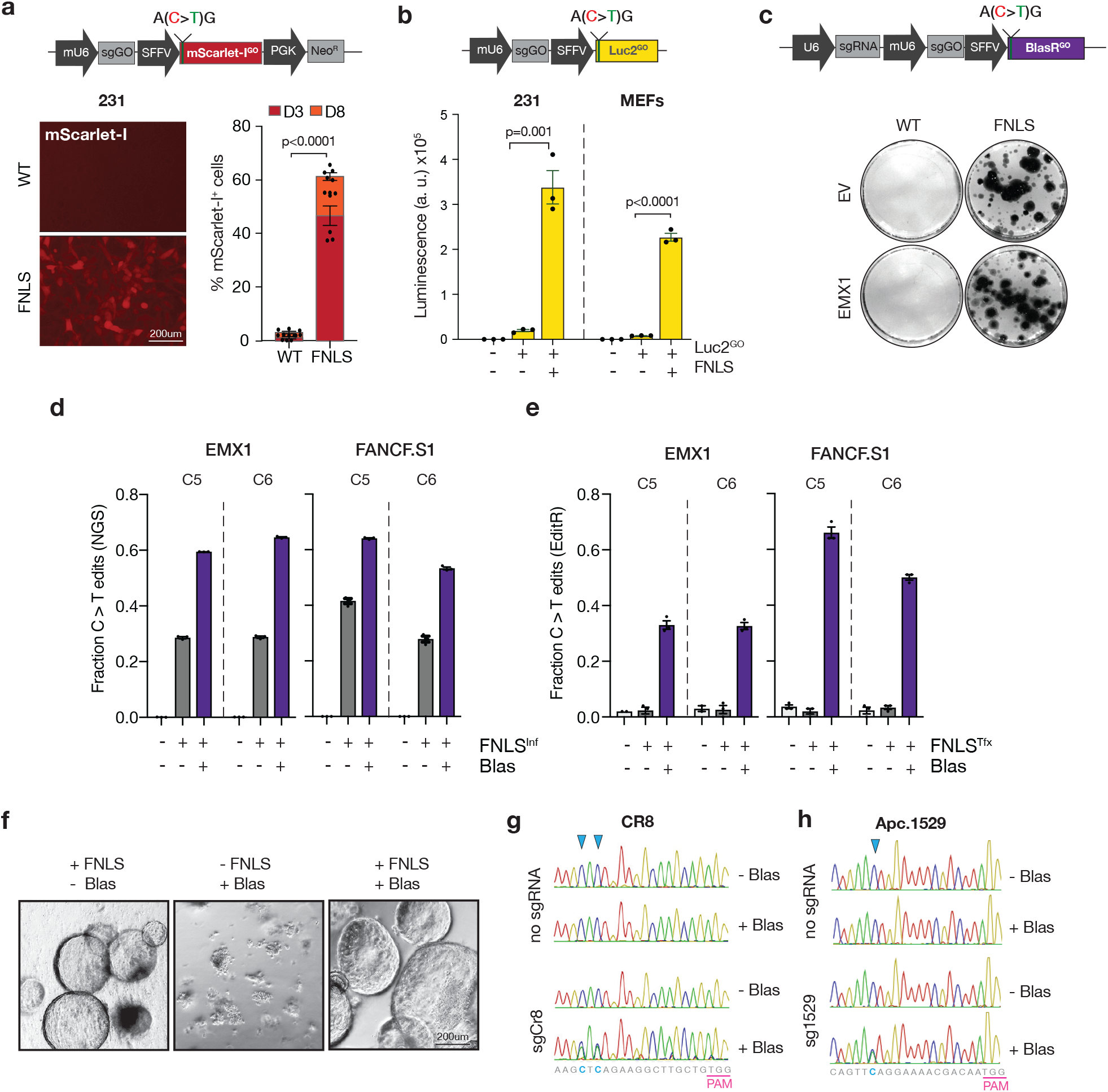
GO is highly modular and can be functionally exploited to enrich base editing activity. **a.** (top) Schematic of an all-in-one mScarlet-I GO reporter with 1) modified 5’ end in the mScarlet-I cDNA to include silent start site ‘ACG’ within 20bp sgGO complementary sequence, 2) sgGO, and 3) neomycin selection gene (Scar^GO^). (bottom, left) Fluorescence microscopy of 231s expressing no editor (WT) or FNLS 3 days after infection with Scar^GO^. (bottom, right) Flow cytometry of 231s expressing no editor (WT) or FNLS 3- and 8-days after infection with ScarGO. **b.** (top) Schematic of an all-in-one-Luciferase GO reporter with 1) a modified 5’ end in the Luciferase2 cDNA to include silent start site ‘ACG’ and 2) sgGO (Luc2^GO^). (bottom) Luminescence in 231s and immortalized MEFs expressing no editor (WT) or FNLS 3 days after infection with and without Luc2^GO^. **c.** (top) Schematic of all-in-one Blasticidin GO reporter with 1) modified 5’ end in the Blasticidin (Blas) resistance gene to include silent start site ‘ACG’ 2) sgGO and, 3) an endogenous targeting sgRNA (Blas^GO^). (bottom) GEMSA staining/colony forming assay of Blas-treated DLD1 cells with FNLS or no editor (WT) infected with Blas^GO^ containing sgRNAs targeting EMX1, FANCF, or no sgRNA (EV). **d.** Deep sequencing analysis of corresponding loci quantifying editing events as a fraction of total reads in DLD1 cells with Blas^GO^(EV, EMX1, FANCF) transduced with and without FNLS or Blas treatment. **e.** EditR quantification of editing events as a fraction of total editing events in DLD1 cells with Blas^GO^ (EV, EMX1, FANCF) transiently transfected with and without FNLS or Blas treatment. **f.** Brightfield images of murine pancreatic organoids with inducible base editor enzyme infected with Blas^GO^ reporters containing sgRNA targeting Apc.1529, CR8, or no guide (EV shown). **g.** Sanger sequencing chromatograms from CR8 loci in pancreatic organoids expressing FNLS and infected with Blas^GO^ (EV or CR8). **h.** Sanger sequencing chromatograms from Apc.1529 loci in pancreatic organoids expressing FNLS and infected with Blas^GO^ (EV or Apc.1529).

### GO enriches endogenous base editing by induction of antibiotic resistance

Isolation of BE in cells with fluorescent reporters requires cell sorting, which may be challenging for some primary cell types or organoid models or due to limited access to specialized equipment. We reasoned that induction of an antibiotic resistance gene would provide a simple and affordable way to select cells with active BE. To test this idea, we designed a silent ‘ACG’ Blasticidin (Blas)-resistance cDNA in an ‘all-in-one’ system containing sgGO under the control of a chimeric U6 promoter, and an empty sgRNA scaffold downstream of the human U6 promoter where user-specific sgRNAs could be integrated (Blas^GO^, Figure 3c). We first cloned two well-characterized sgRNAs (EMX1 and FANCF.S1,) into the Blas^GO^ reporter, and transduced DLD1 cells, which show relatively low BE activity of the cell lines we have examined (Figure 1c). DLD1 cells stably expressing FNLS showed robust outgrowth in the presence of Blas, and Blas-resistant cells had an average 2-fold increase in the frequency of C>T editing at each cytosine of the endogenous targets (Figure 3d, Supplementary Figure 4c).

Stable viral expression of BE enzymes can allow highly efficient gene modification, even in the absence of selection; however, it may induce ongoing off-target effects (17), or cause immune-rejection of cells transplanted into recipient animals (18). Transient transfection of editors bypasses these effects but may also significantly reduce editing efficiency in difficult to transfect cell systems. Indeed, transfection of Blas^GO^ DLD1 cells with FNLS showed minimal evidence of editing in the bulk population (Figure 3e). However, Blas-selected cells showed a dramatic enrichment of editing at the two endogenous targets (EMX1 and FANCF.S1), suggesting the transduction of Blas^GO^ sgRNAs and transient expression of BE enzymes is an effective strategy to isolate edited cells without continual enzyme expression and in otherwise inefficient settings of BE activity.

We have previously used BE in organoids to engineer missense or nonsense mutations in *Pik3ca* or *Apc*, respectively (6, 19). In these cases, modified organoids have a proliferative advantage under restrictive culture conditions and edited populations can be enriched by functional selection. However, isolation of rare BE events that provide no selective advantage in organoids is challenging, as it is difficult to either sort or single-cell-clone primary cells. To test the utility of Blas^GO^ in enriching neutral base editing events in organoids we transduced pancreatic organoids carrying an inducible BE enzyme (6) with a Blas^GO^ vector targeting a non-genic region on mouse chromosome 8 (CR8). Bulk transduced populations had no evidence of targeted editing, while Blas-selected organoids showed approximately 50% edited targets (Figure 3f-g). As a further test we used an established sgRNA targeting *Apc* (Apc.1529;(6)) that creates a non-oncogenic late truncation and provides no growth advantage in organoids). Similar to what we observed with CR8, Blas-selected organoids had more than 50% altered alleles (Figure 3h), dramatically enriching the mutant population and providing an opportunity to engineer and study ‘selection-neutral’ mutations.

### Surrogate Cre-mediated recombination with Cre2^GO^

For modeling disease, BE offers a range of possibilities when combined with existing genetic systems, such as Cre/ LoxP. Linking these two distinct gene targeting events would provide a means to ensure both BE and Cre are active in the same cells. In theory, initiation of Cre expression by GO would trigger LoxP recombination only in cells with active BE, and those cells would have enriched editing at other endogenous genomic sites. To directly test this, we generated MEFs carrying a *R26-LSL-tdTomato* allele (20) that induces red fluorescence following Cre expression. Transduction of this all-in-one Cre^GO^ vector induced tdTomato in more than 70% of MEFs expressing FNLS, but also in nearly 40% of parental cells, suggesting leakiness of Cre expression in the absence of BE (Figure 4a-b, Supplementary Figure 5). We noted that in the first 100 amino acids of Cre there were a number of in frame ATG (Met) and CTG (Leu) codons that could serve as alternate translation start sites. We therefore mutated each CTG>CTC (Leu>Leu) and each ATG>AGT (Met>Ser), creating Cre2^GO^ (Figure 4a). Masking each of the potential alternative start codons had a dramatic impact on Cre2^GO^ specific Cre activity, with FNLS expressing cells showing equivalent levels of editing to Cre^GO^ (>80%), with less than 0.5% tdTomato-positive cells in the absence of FNLS (Figure 4b).

**Figure 4.**
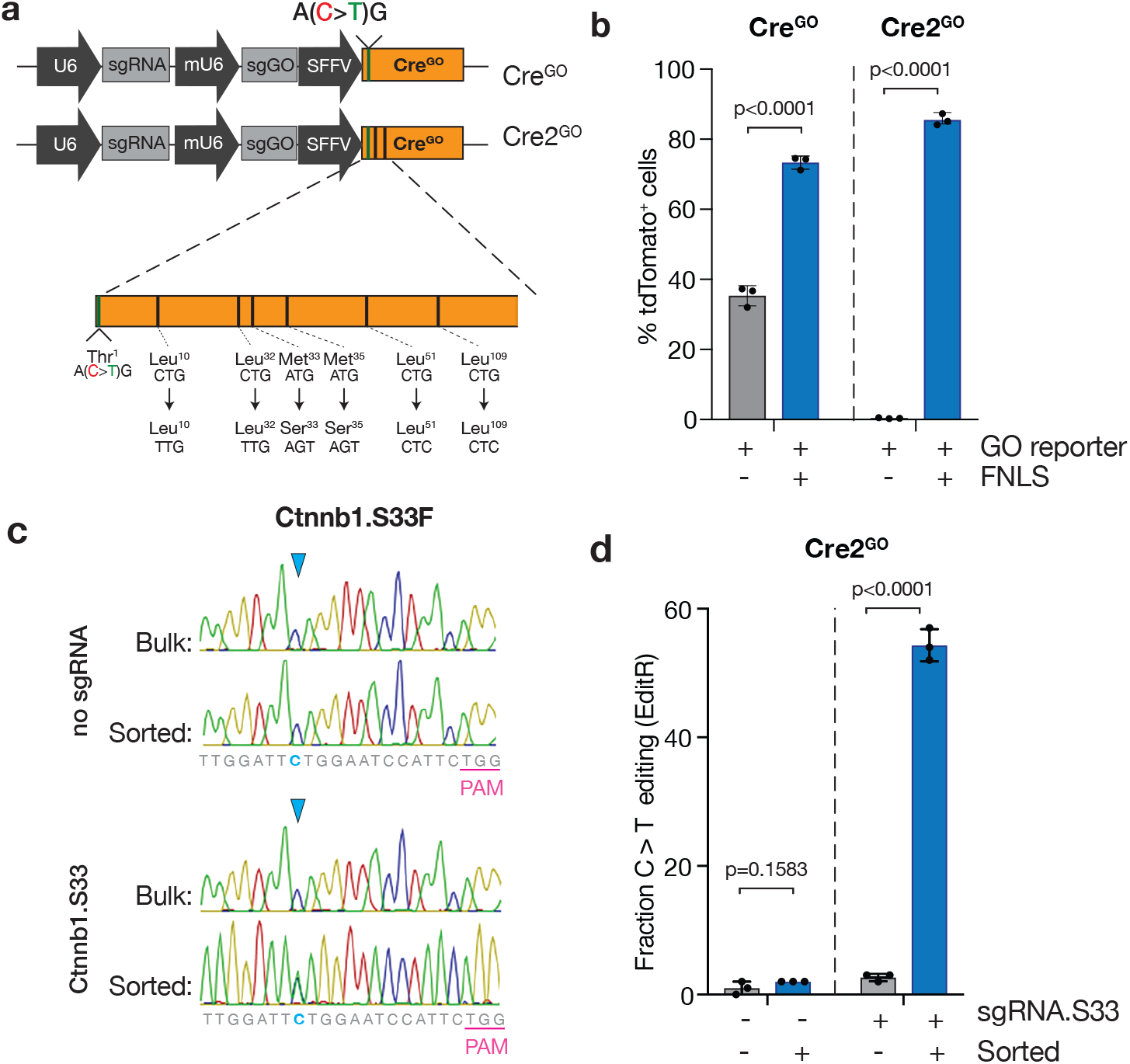
Surrogate Cre-mediated recombination using Cre2^GO^ BE reporter. **a.** (top) Schematic of all-in-one Cre recombinase GO reporter with 1) modified 5’ end in the Cre-recombinase cDNA to include silent start site ‘ACG’, 2) sgGO, and 3) an endogenous targeting sgRNA (Cre^GO^). (bottom) Schematic of modified Cre^GO^ reporter with masked, synonymous alternative start sites (Cre2^GO^). **b.** Flow cytometry of immortalized MEFs containing an *R26-LSL-tdTomato* allele expressing and FNLS or no editor and Cre^GO^ or Cre2^GO^ reporters. FNLS expressing cells contain GFP constitutive marker and were gated on tdTomato+ cells within the GFP^+^ population. WT cells are gated on tdTomato^+^ cells. **c.** *R26-LSL-tdTomato* MEFs expressing FNLS and Cre2^GO^ with sgRNA targeting endogenous Ctnnb1.S33 or no sgRNA (EV) were sorted for tdTomato expression. Sanger sequencing chromatograms are displayed for the bulk and sorted populations at the Ctnnb1.S33 locus. **d.** EditR analysis of sanger sequencing traces quantify editing at the target cytosine for Ctnnb1.S33 in cells from c.

To test the feasibility of enriching endogenous BE by surrogate Cre activity, we generated a Cre2^GO^ reporter with a tandem endogenous gRNA targeting Ser33 of mouse *Ctnnb1*. We simulated restricted BE activity by low MOI transduction with FNLS-P2A-GFP (~10% GFP-positive), and sorted tdTomato-positive cells after 6 days. As observed in GFP^GO^ and Blas^GO^ bulk transduced cells showed very low levels of *Ctnnb1.S33* editing, while tdTomato-positive cells had more than 50% of target cytosines converted (Figure 4c-d). Together, these data show that GO can tether base editing to antibiotic resistance or cellular enzymatic activity, which can be used to functionally enrich endogenous BE events.

## DISCUSSION

Here, we describe a flexible BE reporter system that enables the identification and enrichment of cells and organoids with active cytosine or adenine BE using fluorescence, bioluminescence, antibiotic resistance, or enzymatic markers. In conjunction with the expanding range of optimized BE enzymes, GO vectors can streamline the application of base editing for the generation of targeted genome manipulation in vitro and potentially in vivo.

We first demonstrated the flexibility and quantitative nature of GFP^GO^ to profile a range of new BE variants, identifying the activity and specificity of PAM-flexible and high-fidelity enzymes. Most efforts to characterize PAM specificity and off-target activity use endogenous targets sites that may influence BE activity due to local chromatin features and/or other unknown variables. These tests are critical for defining the real-world effect of specific sgRNAs, but are limiting for direct, quantitative comparisons of BE enzymes. Our experiments revealed cell-type and enzyme-dependent effects on editing efficiency at NGG and NGN PAM sites. While we tested a relatively small panel of BE enzymes and cell lines, GO could be deployed to quickly compare features newly generated enzymes in mammalian cells, optimize editing conditions across different cellular contexts, or broadly classify BE capability of large numbers of cell systems.

A number of recent studies have described BE reporters that induce or inhibit GFP expression by distinct mechanisms. For instance, DOMINO detects base editing by loss of GFP fluorescence (21), while BE-FLARE (22) and TREE (23) generate a fluorescent shift from BFP to GFP after cytidine deamination of codon 66. Finally, Martin et al. established a panel of GFP reporters that restore fluorescence upon C> T base editing at three target codons within GFP (24). Using an adaptable translation initiation approach, we demonstrate the activation of a range of non-GFP reporters. Activating fluorescence markers other than GFP (i.e. mScarlet-I) enables use of GO in systems already utilizing GFP or similarly emitting fluorophores. Alternatively, activating antibiotic resistance with GO facilitates enrichment of endogenous base editing event without the need for flow-based sorting. This is particularly useful in cell systems that are difficult or tedious to sort, such as primary cell types or organoids. It also enables enrichment of endogenous mutations that do not have a natural positive selection, thus increasing the practicality of using BE to model large numbers of disease variants. While we validated this approach using Blas^GO^, in theory the same strategy could be used for other selection markers such as NeoR, PuroR, or HygroR.

In the cases described, GO allows the robust enrichment of endogenous BE events. Yet, in most cases, we were unable to achieve more than 50-80% target C>T editing. In some cases, this was due to non-C>T editing (i.e. C>A or C>G) or the generation of insertions and deletions (indels) at the target site. It is also possible that our current version of GO is a particularly good sgRNA/target combination, increasing the likelihood of reporter activation with even low levels of BE activity. Further, endogenous BE activity is highly dependent on the quality of each individual sgRNA and target site accessibility (25, 26). Given these inherent variables, it will not be possible for any individual reporter to accurately reflect the potency of all sgRNAs. However, the use of sgRNA mismatches or different sgRNA sequences (i.e. GO2) to reduce reporter efficiency may improve enrichment strategies for difficult targets.

BE has enormous potential for *in vivo* modeling of pathogenic mutations but given the variability in editing efficiency across different cell types, it is not clear in which contexts BE will work effectively. Directly tethering BE to activation of Luc2 or Cre will provide a clear readout for in vivo editing efficiency and will enable close integration with established Cre/LoxP systems once the first transgenic BE animal models become available.

GO is a rapid and robust genetic tool that can be used to detect and quantify base editing activity, efficiency, and kinetics. With simple modifications, GO can be adapted to initiate expression of a range of functional reporters, and thus, expands the usability and application of current base editing technologies to include enriching endogenous base editing and triggering a secondary enzymatic activity. We expect GO-based systems will enhance the feasibility of base editing as a go-to genome engineering approach, particularly in difficult-to-manipulate model systems.

## METHODS

### Cloning

GFP^GO^-PGK-Neo lentiviral construct was generated by InFusion assembly (Clontech # 638909) of a custom GFP^ACG^ gBlock cassette (Supplementary Table 1) into AcsI/AgeI-digested SGEN (Addgene #111171) backbone. Ad^GO^-PGK-Neo and Ad^GO2^-PGK-Neo viral constructs were generated by InFusion assembly of custom gBlock cassettes into EcoRI/NsiI-digested GFP^GO^ backbone. GFP^GO2^ PAM variant constructs were generated by amplification of 99mer oligonucleotides with For and Rev primers (Supplementary Table 1) and InFusion assembly into EcoRI-digest GFP^GO^ vector. mUGIS^GO^ and mUGIS^GO2^ were generated by InFusion assembly of IRES and codon-optimized mScarlet-I PCR amplicons into GFP^GO^ backbone. A second version of mUGIS^GO^ was generated incorporating silent mutations in the first 100bp of GFP to eliminate potential CTG alternate start sites. A PGK-Neo cassette was inserted downstream of the mScarlet-I cDNA using by InFusion assembly into SalI-digested mUGIS^GO^ vector. The chimeric mouse U6-sgRNA cassette was generated using a custom gBlock from IDT and cloned into ‘all-in-one’ following PCR amplification and InFusion assembly. Luc2^GO^, Blas^GO^, and Cre^GO^ constructs were generated by InFusion assembly of Luc2/Blas/Cre^GO^ PCR amplicons into digested LRT2B backbone containing mU6-sgRNA cassette and an sgRNA recipient site downstream of the human U6 promoter. Cre2^GO^ was generated by replacing the first ~450bp of Cre with a custom gBlock (IDT) in an AscI/BamHI-digested Cre^GO^ backbone. FNLS-NG was generated using a custom gBlock of codon-optimized sequence incorporating Cas9-NG (13) mutations and assembled using InFusion into digested FNLS backbone (6). FNLS-HiFi was generated using InFusion assembly of 5’ and 3’ FNLS PCR amplicons, incorporating the R691A mutation (16)in the junction of these fragments. FNLS-HF1 and FNLS-HiFi-NG were generated by InFusion assembly of PCR amplicons from FNLS, HF1RA (6), FNLS-HiFi, and FNLS-NG. All sgRNAs were cloned into the BsmBI site of either LRT2B or dual sgRNA recipient vectors (Blas^GO^ and Cre2^GO^) using annealed oligonucleotides (Supplementary Table 2). All other viral vectors used were previously described(6).

### Cell lines

HEK293T (ATCC CRL-3216), MDA-MB-231 (ATCC HTB-26), DLD1 (ATCC CCL-221) and NIH3T3 (ATCC CRL-1658) cell lines were purchased from the ATCC. Stocks were tested for mycoplasma routinely every 6 months and maintained in Dulbecco’s Modified Eagle’s Medium (DMEM, Corning Cat. 10-013-CV) containing 1% Pen/Strep (Corning Cat. 30-002-Cl) and 10% FBS (231 and DLD1) or 10% FCS (3T3s) at 37C with 5% CO2. MEFs from the indicated genotypes were isolated at E13.5 and expanded for 2 passages before transfection. P53KO MEFs were generated by transient transfection of an LCG (Lenti-Cas9-GFP) plasmid containing a p53 gRNA using PEI and selected for p53 disruption by treatment with Nutlin3a at 10ug/ml for 5-10 days. Absence of persistent Cas9 expression was confirmed by Western Blot. MEFs were maintained in DMEM containing 1% Pen/Strep and 10% FBS. Antibiotic selection was performed in complete media as follows: Puromycin (2 days at 1-2ug/ml), Blasticidin (5 days at 5-10ug/ml), Neomycin (3-4 days at 400ug/ml).

### Lentiviral transduction

HEK293T cells were plated at 85-90% confluence in a 6 well and 24h later were transfected with 2.5ug of the vector of interest, 1.25ug of PAX2 and 0.625ug VSVg in 75ul DMEM using a 1:3 DNA:PEI ratio. Cells were washed 24h later in complete media and lentiviral supernatants were collected at 24, 48 and 72h post-transfection. HEK293T cells were cleared out by centrifugation and lentiviral supernatants were diluted 1:4-1:10 prior to target cell infection. Target cells were seeded at 40-60% confluence and incubated with the lentiviral supernatant dilution containing 8ug/ml Polybrene overnight (16h). 24h later we replaced the viral supernatant dilution with complete medium and cells were assayed or subject to selection 48h post-transduction.

### Flow cytometry

We used a Thermo Fisher 2018 Attune NxT flow cytometer. Cells were trypsinized at the indicated time points and resuspended in 300ul of complete medium. Flow cytometry assays were carried out in round-bottom 96 well plates and data were acquired at a flow rate of 500ul/min. At least 25,000 events from the single cell population gating were recorded, and all experiments were performed in triplicates coming from independent transductions. Weill Cornell Flow cytometry core performed sorting using a BD FACS Aria II Cell Sorter.

### Luciferase Activity

Cells were cultured in 96 well flat-bottom plates in 200uL of complete media. Cells were stimulated by Dual-Glo Luciferase Assay System (Promega) and luminescence measured on 96-well plate reader.

### Organoid culture, transfection, and transduction

Murine small intestine organoids from the indicated genotypes were isolated and maintained as previously described (27). Isolation of murine pancreatic ductal organoids was done modifying previously described protocol (28). Briefly, pancreas was minced and washed in Hanks’s Balanced Salt Solution (Corning), and then incubated for 30 min at 37°C with Collagenase V to release the ducts. After washing twice with DMEM/10% FBS media, ducts were resuspended in basal media [Advanced DMEM/F12 (Corning) containing 1% penicillin/streptomycin, 1% glutamine, 1.25 mM N-acetylcysteine (Sigma Aldrich A9165-SG) and B27 Supplement (Gibco)], and mixed 1:10 with factor reduced (GFR) Matrigel (BD Biosciences). Forty microliters of the resuspension was plated per well in a 48-well plate and placed in a 37°C incubator to polymerize for 10 minutes. To culture ductal pancreatic organoids the basal media described above was supplemented with 10 nM Gastrin (Sigma), 50 ng/ml EGF (Peprotech), 10% RSPO1-conditioned media, 100 ng/ml Noggin (Peprotech), 100 ng/ml FGF10 (Peprotech) and 10 mM Nicotinamide (Sigma). Note: Culture freshly isolated organoids in pancreatic organoid media (POM) containing 10 mM Rock inhibitor (Y2732) during 72-48 h. For subculture and maintenance, media were changed on organoids every two days and they were passaged 1:3 every 5 days. To passage, the growth media was removed and the Matrigel was resuspended in cold basal media and transferred to a 15-mL Falcon tube. Organoids were mechanically disassociated using a P1000 and pipetting 40 times. Five milliliters of cold PBS were added to the tube and cells were then centrifuged at 1,200 rpm for 5 minutes and the supernatant was aspirated. Cells were then resuspended in GFR Matrigel and replated as above. For freezing, after spinning the cells were resuspended in complete containing 10% FBS and 10% DMSO and stored in liquid nitrogen indefinitely. Organoids were transfected or transduced as previously described (29). Antibiotic selection was performed in complete media as follows: Blasticidin S (7 days at 5ug/ml), Neomycin (7 days at 100ug/ml).

### PCR amplification for sequencing

Target genomic regions of interest were amplified by PCR. PCR was performed with Herculase II Fusion DNA polymerase (Agilent Technologies, Palo Alto, CA, USA, #600675) according to the manufacturer’s instructions using 200 ng of genomic DNA as a template and under the following PCR conditions: 95C × 2 min, 95C - 0:20, 58C - 0:20, 72C - 0:30 × 34 cycles, 72C × 3 min (6). PCR produces were purified using QIAquick PCR Purification Kit (Qiagen Cat. 28106).

### Sequencing

Sanger sequencing reactions were conducted at Eton Bioscience (Union, NJ, USA). Sanger sequencing chromatograms were visualized using Geneious version 10.2.6 created by Biomatters and analyzed using EditR (30). Targeted amplicon library preparation and NGS sequencing (MiSeq; 2 x 250bp) were performed at GENEWIZ, Inc. (South Plainfield, NJ, USA) and analyzed using CRISPResso2 (31).

### Statistical Analysis

All statistical tests used throughout the manuscript are indicated in the appropriate figure legends. In general, to compare two conditions, a standard two-tailed unpaired t test was used, assuming variance between samples. In most cases, analyses were performed with one-way or two-way ANOVA, with Tukey’s correction for multiple comparisons. Unless otherwise stated, each replicate represents an independent cell transfection or independently transduced cell line. Results of all statistical tests are available in Supplemental Table 3.

### Ethics approval and consent to participate

This study did not involve human subjects. All animal experiments were approved by the Weill Cornell Medicine Institutional Animal Care and Use Committee (IACUC) under protocol 2014-0038.

### Availability of data and materials

Raw MiSeq fastq files have been deposited in the sequence read archive (SRA) under accession PRJNA588416. All plasmids described will be made available at the non-profit repository Addgene.

## Acknowledgements

We thank members of the Dow lab for advice and comments on preparation of the manuscript. This work was supported by a project grant from the NIH/NCI (R01CA229773-01A1). MPZ is supported in part by National Cancer Institute (NCI) Grant NIH T32 CA203702. The content is solely the responsibility of the authors and does not necessarily represent the official views of the NIH.

## Authors’ contributions

AK and MF designed and performed experiments, analyzed data and wrote the paper. JZ, BD, MPZ, and SG performed experiments and/or analyzed data. LED designed and supervised experiments, analyzed data, and wrote the paper.

---

**Supplementary Figure 1.**
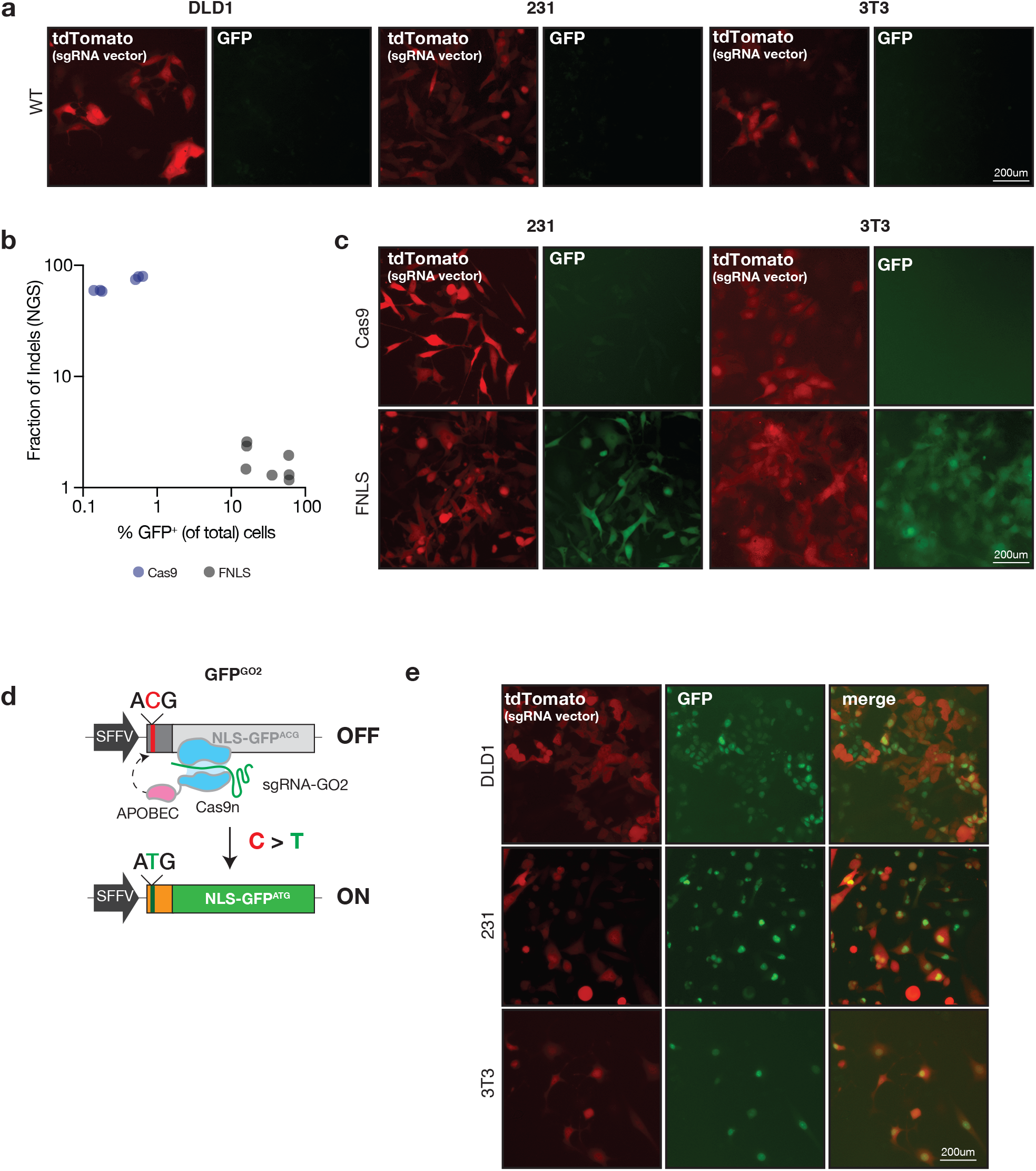
**a.** Fluorescence microscopy of DLD1, 231 and 3T3 parental cells infected with GFP^GO^ and sgGO-tdTomato without editor. **b.** Correlation of targeted deep sequencing insertion/deletion (indel) editing events as a fraction of total reads relative to GFP^+^ (not gated on tdTomato) cells. 231, DLD1, and 3T3 cells containing GFP^GO^, sgGO-tdTomato, and Cas9 or FNLS were analyzed 3 days after sgGO transduction. **c**. Fluorescence imaging of 231 and 3T3 cells infected with GFP^GO^, sgGO-tdTomato, and Cas9 or FNLS 3 days after transduction with sgGO. **d.** GFP^GO2^ schematic designed to introduce an NLS-tagged GFP^GO^ reporter and corresponding targeting sgRNA (sgGO2). **e.** Fluorescence imaging of DLD1, 231, and 3T3 cells infected with GFP^GO2^, sgGO2-tdTomato, and FNLS, 3 days after transduction with sgGO.

**Supplementary Figure 2.**
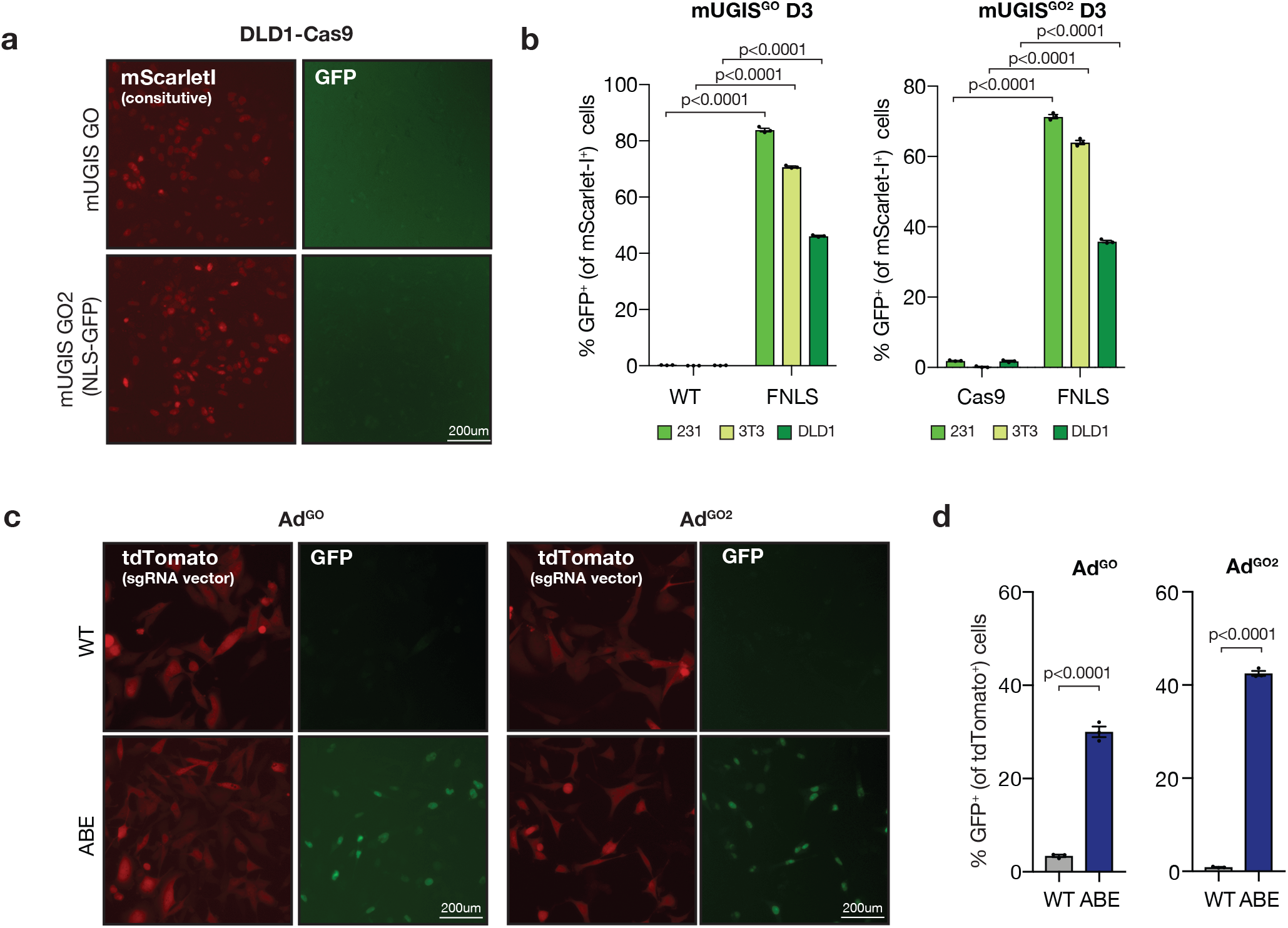
**a.** Fluorescence microscopy of DLD1 cells expressing Cas9 day 3 after infection with mUGIS^GO^ or mUGIS^GO2^. **b**. Flow cytometric analysis of DLD1, 231, and 3T3 cells expressing FNLS or (left) no editor (WT) day 3 after infection with mUGIS^GO^ or (right) Cas9 day 3 after infection with mUGIS^GO2^. GFP^+^ cells were gated within the mScarlet-I^+^ population. **c.** Fluorescence microscopy of 231 cells infected with Ad^GO^ or Ad^GO2^ and FNLS or no editor 3 days after transduction with corresponding sgRNA, sgAd^GO^ or sg Ad^GO2^, respectively. **d.** Flow cytometric analysis of 231 cells infected with Ad^GO^ or Ad^GO2^ and FNLS or no editor, 3 days after transduction with corresponding sgRNA, sgAd^GO^ or sg Ad^GO2^, respectively.

**Supplementary Figure 3.**
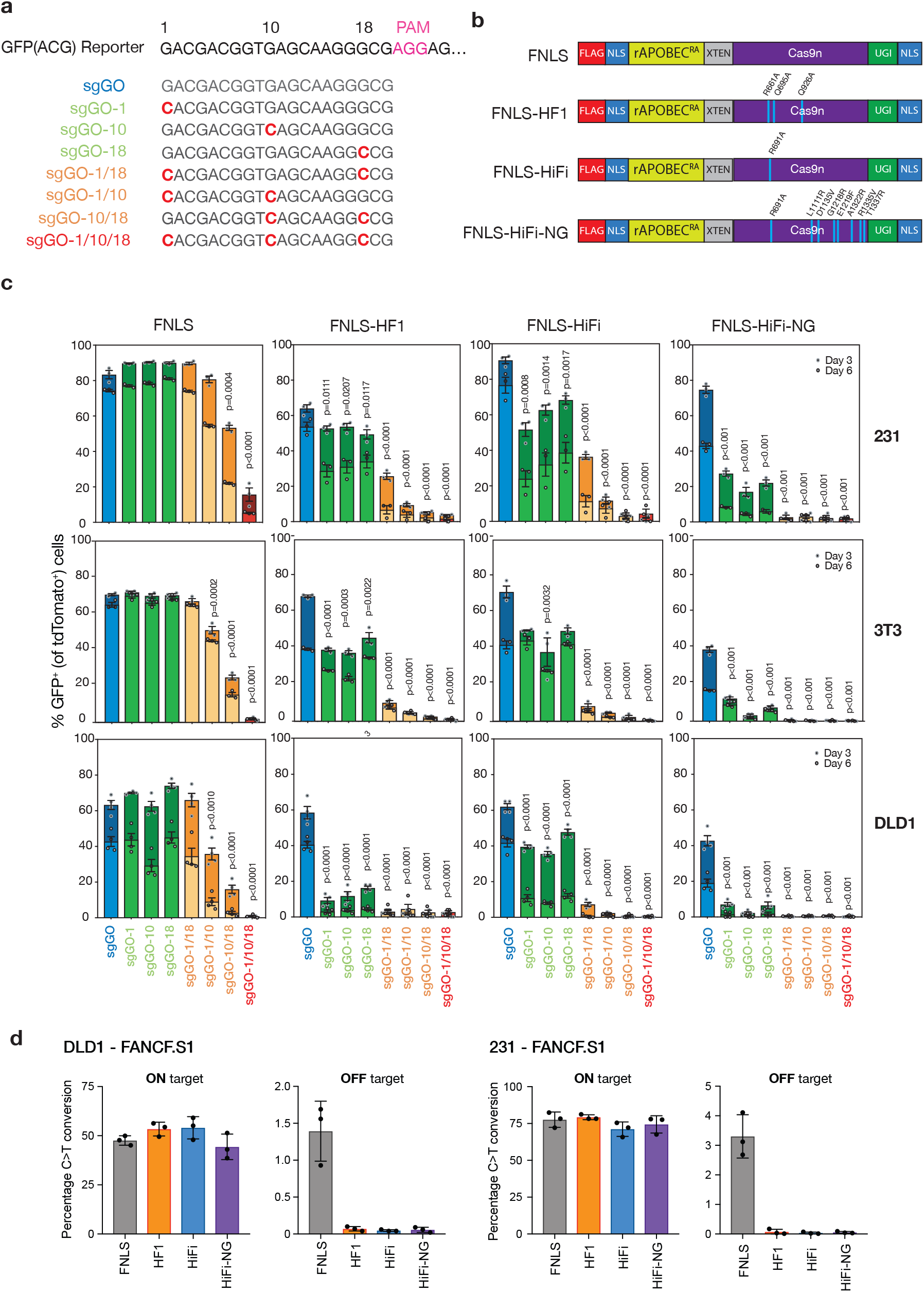
**a.** A panel of sgRNAs targeting the 5’sequence of the GFP^ACG^ cDNA modified from sgGO were designed to include 1 to 3 mismatches across its complementary sequence. **b.** Schematic of high fidelity BE enzymes including novel HIFI-NG enzyme created by combining mutations indicated to improve fidelity, and/or PAM flexibility. **c**. 231, DLD1, and 3T3 cells expressing GFP^GO^ and then panel of high-fidelity BE enzymes after 3 and 6 days post transduction with the panel of mismatch sgGOs. GFP^+^ cells were gated on tdTomato positivity. **d.** DLD1 and 231 cells expressing the panel of high fidelity BE enzymes were infected with an sgRNA targeting FANCF and a known off target locus with two mismatches at position 5 and 9 within the complementary guide sequence. Deep sequencing analysis quantifies C>T editing events as a fraction of total reads at the on target (FANCF) and off-target locus.

**Supplementary Figure 4.**
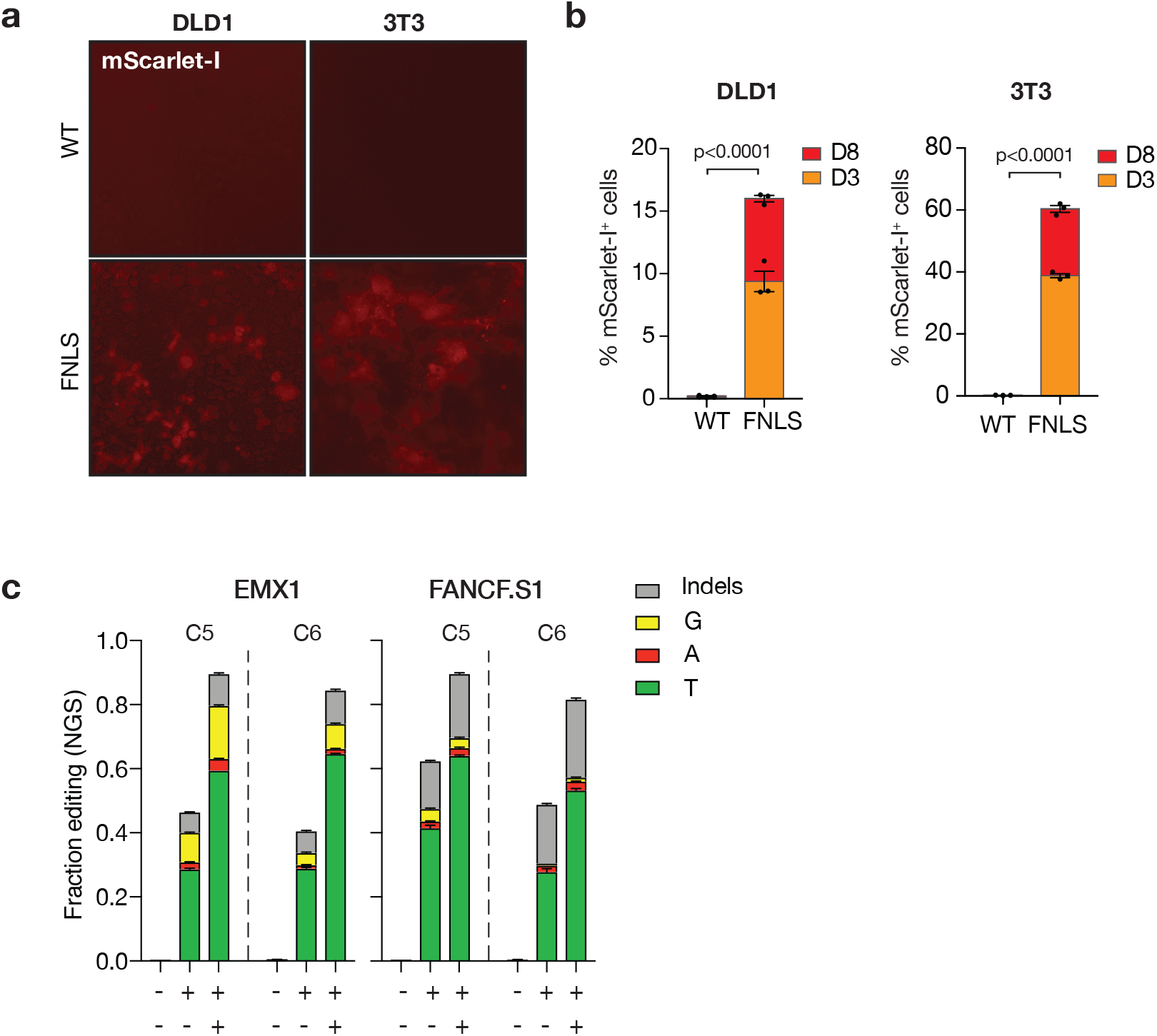
**a.** Fluorescence microscopy images of DLD1 and 3T3 cells expressing FNLS 6 days after infection with Scar^GO^. **b.** Flow cytometry analysis of DLD1 and 3T3 cells 3 and 8 days after infection with Scar^GO^ with and without FNLS. **c.** Deep sequencing analysis of corresponding loci quantifying all possible editing events (including C to T, redundant with Figure 3d) as a fraction of total reads in DLD1 cells with Blas^GO^ (EV, EMX1, FANCF, CTNNB1) transduced with and without FNLS or Blas treatment.

**Supplementary Figure 5.**
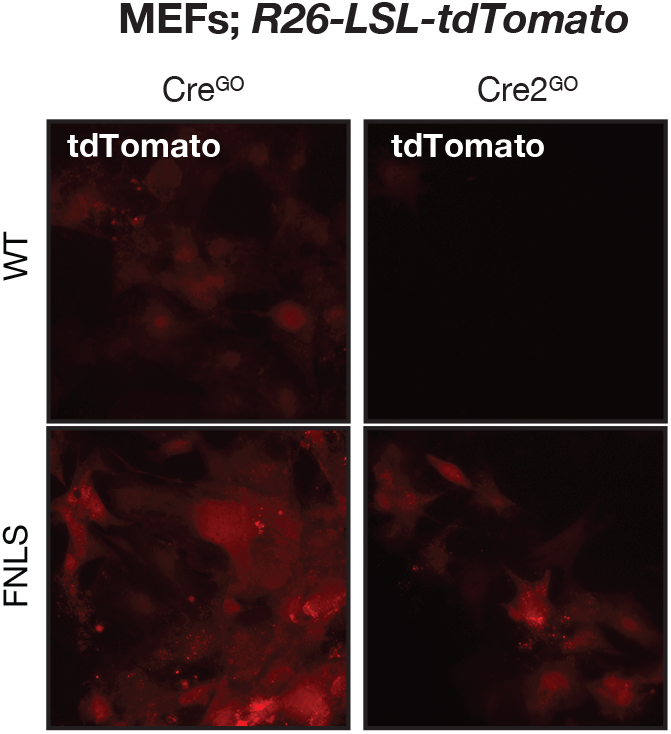
**a.** Fluorescence microscopy images of *R26-LSL-tdTomato* containing immortalized MEF cells expressing FNLS or no editor (WT) 3 days after infection with Cre^GO^ or Cre2^GO^.

